# A simple microbiome in the European common cuttlefish, *Sepia officinalis*

**DOI:** 10.1101/440677

**Authors:** Holly L. Lutz, S. Tabita Ramírez-Puebla, Lisa Abbo, Amber Durand, Cathleen Schlundt, Neil Gottel, Alexandra K. Sjaarda, Roger T. Hanlon, Jack A. Gilbert, Jessica L. Mark Welch

**Affiliations:** Department of Pediatrics, University of California San Diego, La Jolla, California, USA; Scripps Institution for Oceanography, UCSD, La Jolla, California, USA; Integrative Research Center, Field Museum of Natural History, Chicago, Illinois, USA; Marine Biological Laboratory, Woods Hole, MA 02543; Biological Sciences Division, University of Chicago, Chicago, IL 60637

**Keywords:** microbiome, fluorescence *in situ* hybridization, Cephalopoda, Vibrionaceae, Piscirickettsiaceae, enrofloxacin

## Abstract

The European common cuttlefish, *Sepia officinali*s, is used extensively in biological and biomedical research yet its microbiome remains poorly characterized. We analyzed the microbiota of the digestive tract, gills, and skin in mariculture-raised *S. officinalis* using a combination of 16S rRNA amplicon sequencing, qPCR and fluorescence spectral imaging. Sequencing revealed a highly simplified microbiota consisting largely of two single bacterial amplicon sequence variants (ASVs) of Vibrionaceae and Piscirickettsiaceae. The esophagus was dominated by a single ASV of the genus *Vibrio*. Imaging revealed bacteria in the family Vibrionaceae distributed in a discrete layer that lines the esophagus. This *Vibrio* was also the primary ASV found in the microbiota of the stomach, cecum, and intestine, but occurred at lower abundance as determined by qPCR and was found only scattered in the lumen rather than in a discrete layer via imaging analysis. Treatment of animals with the commonly-used antibiotic enrofloxacin led to a nearly 80% reduction of the dominant *Vibrio* ASV in the esophagus but did not significantly alter the relative abundance of bacteria overall between treated versus control animals. Data from the gills was dominated by a single ASV in the family Piscirickettsiaceae, which imaging visualized as small clusters of cells. We conclude that bacteria belonging to the Gammaproteobacteria are the major symbionts of the cuttlefish *Sepia officinalis* cultured from eggs in captivity, and that the esophagus and gills are major colonization sites.

**IMPORTANCE:** Microbes can play critical roles in the physiology of their animal hosts, as evidenced in cephalopods by the role of *Vibrio (Aliivibrio) fischeri* in the light organ of the bobtail squid and the role of Alpha- and Gammaproteobacteria in the reproductive system and egg defense in a variety of cephalopods. We sampled the cuttlefish microbiome throughout the digestive tract, gills, and skin and found dense colonization of an unexpected site, the esophagus, by a microbe of the genus *Vibrio*, as well as colonization of gills by Piscirickettsiaceae. This finding expands the range of organisms and body sites known to be associated with *Vibrio* and is of potential significance for understanding host-symbiont associations as well as for understanding and maintaining the health of cephalopods in mariculture.

## 1. INTRODUCTION

Symbiotic associations between invertebrates and bacteria are common. Among cephalopods, the most intensely studied association is the colonization of the light organ of the bobtail squid *Euprymna scolopes* by the bioluminescent bacterium *Vibrio (Aliivibrio) fischeri* in a highly specific symbiosis (1). A more diverse but still characteristic set of bacteria colonize the accessory nidamental gland from which they are secreted into the egg jelly coat and likely protect the eggs from fungal and bacterial attack (2). The accessory nidamental gland and egg cases of the squid *Doryteuthis* (*Loligo) pealeii* and the Chilean octopus (*Octopus mimus*) have also been reported to contain Alphaproteobacteria and Gammaproteobacteria (3, 4). These associations indicate that bacteria can play a key role in the physiology of cephalopods.

*Sepia officinali*s, the European common cuttlefish (hereafter cuttlefish), is used extensively in biological and biomedical research (5–7) and is a model organism for the study of rapid adaptive camouflage (8–11). Cuttlefish are also widely represented among zoos and aquaria, and play an important role in educating the public about cephalopod biology and life history (12). Little is known about the association of bacterial symbionts with cuttlefish, and whether such associations may play a role in the health or behavior of these animals. Understanding the importance, or lack thereof, of the cuttlefish microbiome will not only shed light on the basic biology of this model organism but will also have important implications for future husbandry practices and research design.

Using a combination of 16S rRNA amplicon sequencing, fluorescence *in situ* hybridization (FISH), and quantitative PCR (qPCR), we characterized the gastrointestinal tract (GI), gill, skin, and fecal microbiota of the common cuttlefish in wild-bred, captive-raised animals (5) housed at the Marine Biological Laboratory (Woods Hole, MA). We observed a highly simplified microbiome dominated by Vibrionaceae in the gastrointestinal tract and Piscirickettsiaceae in the gills. We treated a subset of cuttlefish with antibiotic enrofloxacin, commonly used among aquaria veterinarians, and found both ASVs to remain dominant in esophagus and gill microbiota, suggesting they are resilient to this antibiotic. The simplicity of this system makes it a promising model for further exploration of the factors driving host-symbiont associations in marine invertebrates.

## 2. RESULTS

### 2.1 Two taxa dominate the *S. officinalis* microbiome

We sampled 27 healthy adult cuttlefish (*Sepia officinalis*) from the mariculture laboratory at Marine Biological Laboratory (Woods Hole, MA). The study comprised two time periods. The first (20-21 June 2017) was a pilot survey in which three individuals were sampled. The second (25 September −10 October 2017) was an experiment involving 24 individuals, of which 16 were exposed to repeated doses of the antibiotic enrofloxacin and 8 individuals served as untreated controls. 16S rRNA amplicon sequencing of the GI tract, gills, and skin of all 27 animals revealed a highly simplified microbiota dominated by bacterial amplicon sequence variants (ASVs) in the Vibrionaceae and Piscirickettsiaceae families, regardless of treatment with enrofloxacin.

In particular, results showed a consistent and highly simplified microbiota in the esophagus (Figure 1; Table 1). A single ASV in the genus *Vibrio* (referred to as ASV1 in subsequent figures and tables) made up the majority of the 16S rRNA sequence data from the esophagus of the three pilot investigation individuals (mean 92% ± 10%) and of 24 individuals sampled four months later (control group mean 100% ± 1%; treatment group mean 94% ± 10%). Thus, this ASV represents a dominant constituent of the esophagus microbiota stably over two time periods in the study and after exposure to antibiotic treatment with enrofloxacin. Another ASV of the related genus *Photobacterium* (Vibrionaceae) (referred to as ASV2 in Table 1) was present in the esophagus community in the pilot investigation animals (mean 7% ± 10%). Combined, the two Vibrionaceae ASVs (ASV1 and ASV2) in the three pilot animals constituted >99% of the esophagus community.

**Figure 1.**
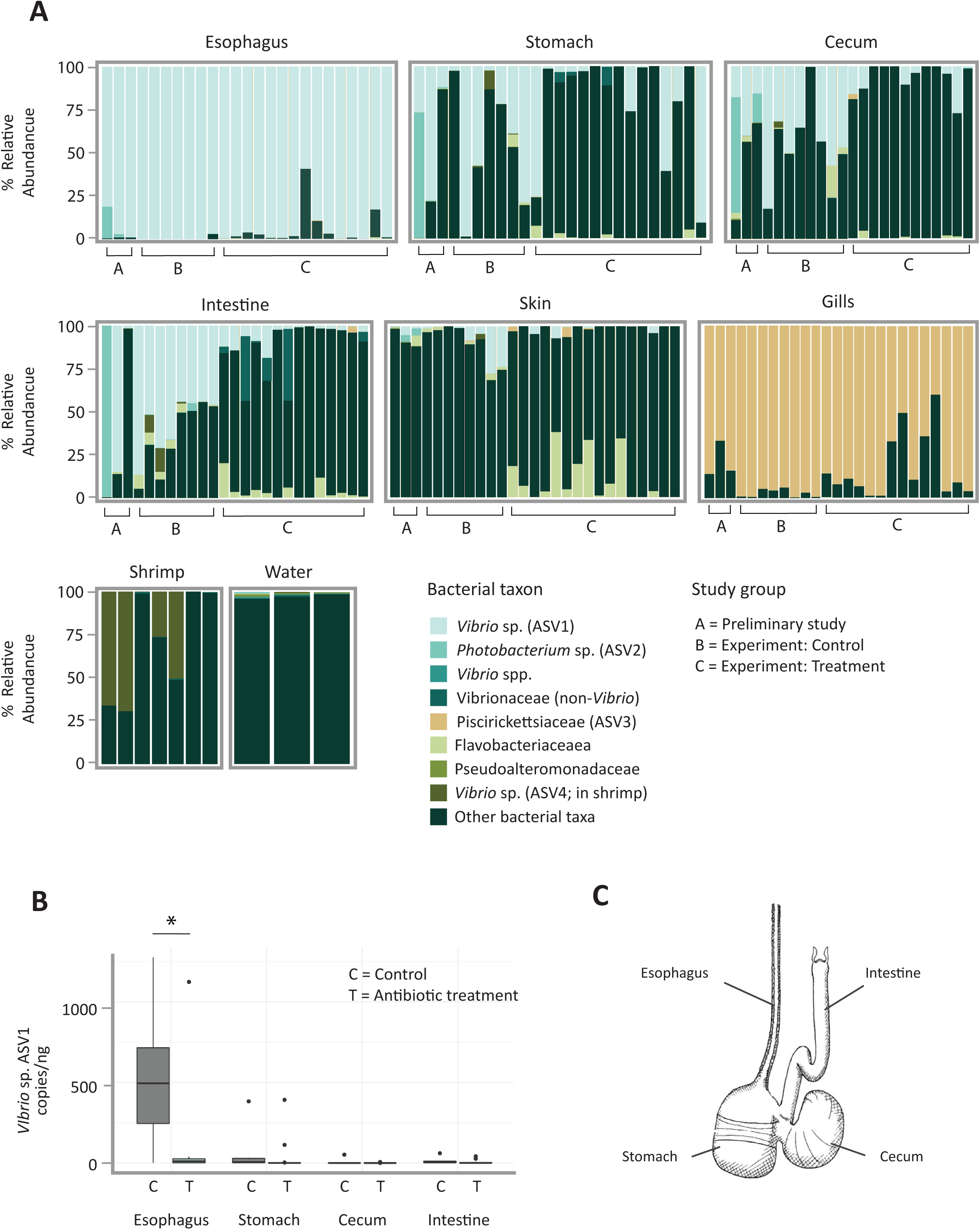
A single *Vibrio* taxon dominates the esophagus, and a single Piscirickettsiaceae taxon dominates the gills of the European common cuttlefish in captivity. (A) Relative abundance of top bacterial taxa found among cuttlefish organs. Shrimp for feeding and seawater from holding tanks are also shown. ASVs are labeled according to the finest level of taxonomic resolution provided by the Greengenes database; bacteria not included in the top 8 taxa are pooled as “other”. Bars correspond to individual 16S rRNA sequence libraries from the pilot investigation animals (labeled “A”), experimental animals in the control category (labeled “B”), and experimental animals in the antibiotic treatment category (labeled “C”); only libraries with >1000 read depth are shown. (B) Quantity of *Vibrio* cells per nanogram of DNA measured using *Vibrio*-specific primers (567F and 680R (43)). Asterisk indicates significant difference between organs (p < 0.05, Welch two sample t-test). (C) Anatomical depiction of the gastrointestinal tract of *S. officinalis*, illustration modified from Plate XI of Tompsett (44).

**Table 1.** Relative abundance of three most abundant ASVs found among two sampling periods of the European common cuttlefish. Averages correspond to relative abundance of individual ASVs from each anatomical site and period. Period 1 consisted of three individuals, Period 2 consisted of eight individuals.

| Anatomical Site | ASV1 |  |  | ASV2 |  |  | ASV3 |  |  |
| --- | --- | --- | --- | --- | --- | --- | --- | --- | --- |
|  | Vibrionaceae, *Vibrio* sp. |  |  | Vibrionaceae, *Photobacterium* sp. |  |  | Piscirickettsiaceae, *unknown* |  |  |
|  | Pilot | Control | Antibiotic | Pilot | Control | Antibiotic | Pilot | Control | Treatment |
| Esophagus | 0.92 ±<br>0.10 | 1.00 ±<br>0.01 | 0.94 ±<br>0.10 | 0.07 ±<br>0.10 | 0 ± 0 | 0 ± 0 | 0 ± 0 | 0 ± 0 | 0 ± 0 |
| Stomach | 0.39 ±<br>0.34 | 0.43 ±<br>0.37 | 0.19 ±<br>0.31 | 0.25 ±<br>0.42 | 0 ± 0 | 0 ± 0 | 0 ± 0 | 0 ± 0 | 0 ± 0 |
| Cecum | 0.24 ±<br>0.14 | 0.44 ±<br>0.23 | 0.06 ±<br>0.09 | 0.28±0.35 | 0 ± 0 | 0 ± 0 | 0 ± 0 | 0 ± 0 | 0 ± 0 |
| Intestine | 0.22 ±<br>0.42 | 0.57 ±<br>0.16 | 0.06 ±<br>0.06 | 0.50 ±<br>0.57 | 0.01 ±<br>0.01 | 0 ± 0 | 0 ± 0 | 0 ± 0 | 0 ± 0 |
| Gills | 0 ± 0 | 0 ± 0 | 0 ± 0 | 0 ± 0 | 0.01 ±<br>0.01 | 0 ± 0 | 0.79 ±<br>0.11 | 0.97 ±<br>0.03 | 0.82 ±<br>0.19 |
| Skin | 0.02 ±<br>0.03 | 0.08 ±<br>0.11 | 0.01 ±<br>0.02 | 0.03 ±<br>0.02 | 0 ± 0 | 0 ± 0 | 0 ± 0 | 0 ± 0 | 0.01 ±<br>0.02 |

The major *Vibrio* ASV1 was also a major constituent of downstream sites in the GI tract, although present at lower abundance measured both as relative abundance in 16S sequencing data and by quantitative PCR (qPCR) (Figure 1; Table 1). qPCR revealed a high abundance of *Vibrio* cell copies in the esophagus (average 520.4 ± 410 copies/ng of total DNA including host DNA) relative to more distal portions of the GI tract that included stomach, cecum, and intestine (combined average 25.6 ± 77.8 copies/ng) (p < 0.005, χ^2^ = 16.5, df =3 Kruskal-Wallis) (Fig. 1B). Comparison of qPCR measures of ASVs in the genus *Vibrio* between the esophagus of treatment and control animals revealed a striking and significant decrease of nearly 80% in the quantity (p < 0.02, df = 10.8, Welch two sample t-test). We did not observe a significant difference in *Vibrio* ASV1 quantity between other organs of the digestive tract with antibiotic treatment (Fig. 1B). Relative abundance of *Vibrio* ASV1 in the esophagus and stomach also did not differ significantly between antibiotic and control groups (Welch two sample t-test, p > 0.10), but did differ significantly among the cecum (Welch two sample t-test, p = 0.00235) and intestine (Welch two sample t-test, p = 1.99e-05) of treatment versus control groups. Analysis of GI tracts (esophagus, stomach, cecum, and intestine samples combined) between treated versus control animals from the second period of study revealed significant differences in weighted UniFrac β-diversity between the two groups (PERMANOVA Pr(>F) = 0.006, F = 5.63, df = 1), despite the *Vibrio* ASV1 remaining dominant in most organs. These differences in measured relative abundance and β-diversity may result from stochastic variation in low-abundance sequences, as the non-*Vibrio* portion of the 16S rRNA sequence data from the GI tract consisted of an assortment of taxa that varied between individuals or between the two time points of the study and thus suggested transient organisms rather than stable microbial colonization.

Samples from gills were dominated by a single highly abundant ASV in the family Piscirickettsiaceae (referred to as ASV3 in subsequent figures and tables), which made up an average relative abundance of 96.9% ± 2.5% in the gills. In samples from other body sites this ASV3 was detected only sporadically and at low relative abundance (mean 0.2%, range 0 to 5.8%) (Fig. 1A). An additional Vibrio ASV, ASV4, was a major constituent of the microbiota of the shrimp used as food for the cuttlefish and was also detectable in some samples from stomach, cecum, and intestine (Figure 1A). Skin samples did not exhibit much similarity to GI tract or gills with respect to microbiome composition, with most common ASVs found in other anatomical sites comprising < 20% (mean 17.2% ± 2.9%) relative abundance of the microbiome (Table S1).

In addition to surveying internal organs, we collected fecal samples from the 24 animals from the second time period of our study. These samples were collected daily for each individual throughout the course of the antibiotic treatment experiment (see methods). Comparison of weighted UniFrac dissimilarity of fecal samples from experimental animals revealed significant differences in beta-diversity between cuttlefish fecal samples, seawater, and shrimp (PERMANOVA Pr(>F) = 0.001, F = 5.26, df = 2) upon which the animals were fed. These results, paired with the differences we observed in compositional relative abundance between organs, seawater, and shrimp provide additional support for our finding that bacterial communities associated with cuttlefish differ from those found in their seawater environment and food source (Fig. 2).

**Figure 2.**
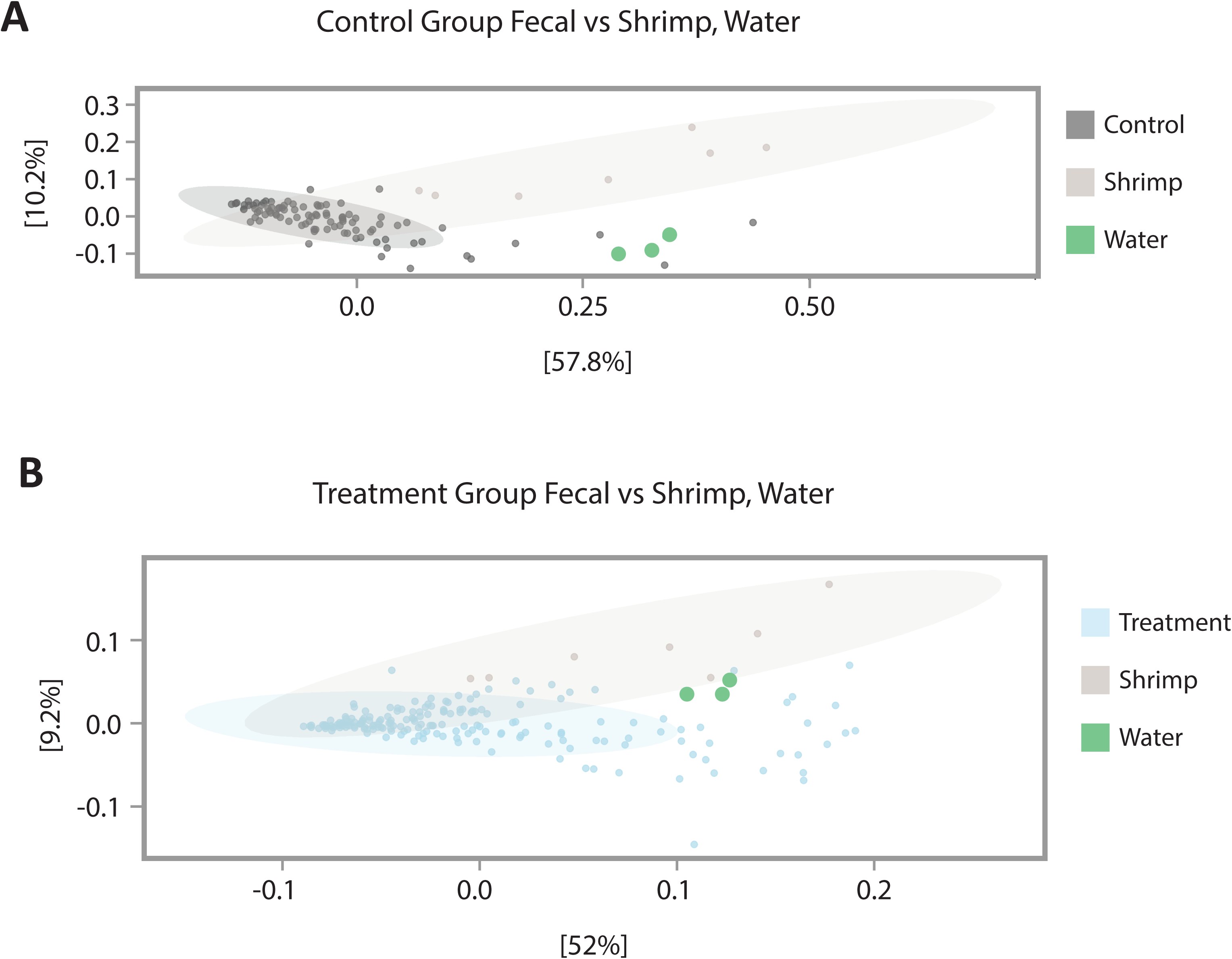
Principal Coordinates Analysis (PCoA) of weighted UniFrac β-diveristy comparing shrimp, tank water, and (A) fecal samples of treatment cuttlefish and (B) fecal samples of control cuttlefish. Fecal samples from control animals (B) show a more tightly clustered pattern than do fecal samples from animals treated with enrofloxacin (A).

### 2.2 Imaging shows spatial structure of the microbiota in cuttlefish esophagus and scattered distribution elsewhere in the gastrointestinal tract

Fluorescence *in situ* hybridization (FISH) revealed a striking organization of bacteria distributed in a layer lining the interior of the convoluted esophagus of cuttlefish (Fig. 3A-C). Hybridization with the near-universal Eub338 probe showed bacteria in high density in a layer ∼20-40 μm thick at the border between host tissue and lumen. Staining with fluorophore-conjugated wheat germ agglutinin revealed a mucus layer that covered the epithelium and generally enclosed the bacteria (Fig. 3). To verify the identity of these bacteria we employed a nested probe set including Eub338 as well as probes for Alphaproteobacteria, Gammaproteobacteria, and probes we designed specifically for Vibrionaceae (Vib1749 and Vib2300, Table 2). Bacterial cells imaged in the esophagus showed signal from all probes expected to hybridize with Vibrionaceae, suggesting that the bacteria observed in this organ are a near-monoculture of this taxon (Fig. 4B-E). A probe targeted to Alphaproteobacteria was included in the FISH as a negative control and, as expected, did not hybridize with the cells (Fig. 4F). As an additional control to detect non-specific binding of probes, we performed an independent FISH with a set of probes labeled with the same fluorophores as the experimental probe set but conjugated to oligonucleotides not expected to hybridize with the cuttlefish microbiota (Table 2). No signal from this non-target probe set was detected (Fig. 4G-H) supporting the interpretation that the signal observed in the esophagus results from a specific interaction of the Vibrionaceae-targeted oligonucleotides with the visualized bacteria.

**Figure 3.**
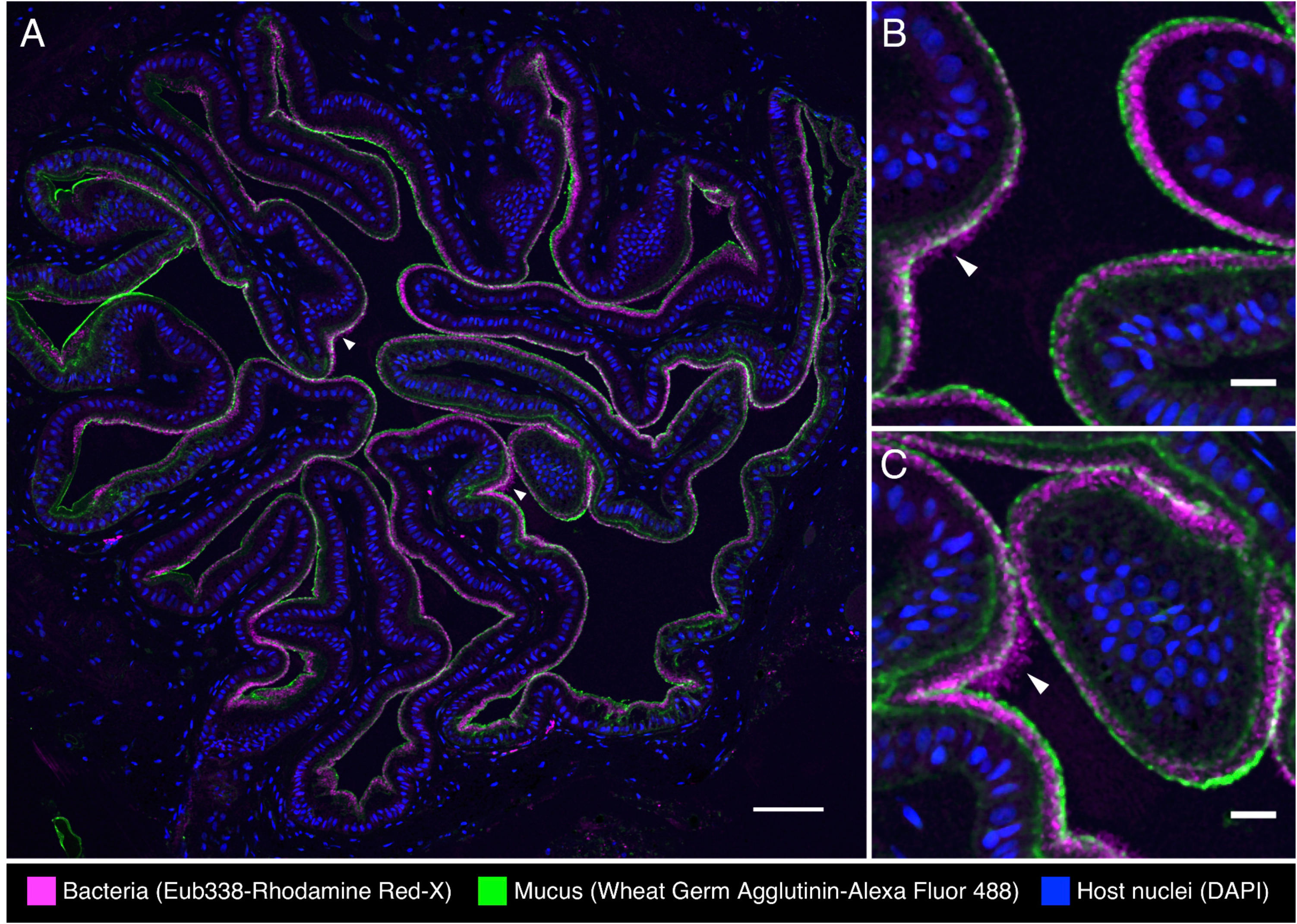
Spatial organization of bacteria in the esophagus of the European common cuttlefish, *S. officinalis*. The image shown is a cross-section of esophagus that was embedded in methacrylate, sectioned, and subjected to fluorescence *in situ* hybridization (FISH). (A) Bacteria (magenta) lining the interior of the esophagus in association with the mucus layer (wheat germ agglutinin staining, green). (B) and (C) are enlarged images of the regions marked with arrowheads in (A) where bacteria extend past the edge of the mucus layer. Blue: Host nuclei. Scale bar =100 µm (A); 20 µm (B) and (C).

**Figure 4.**
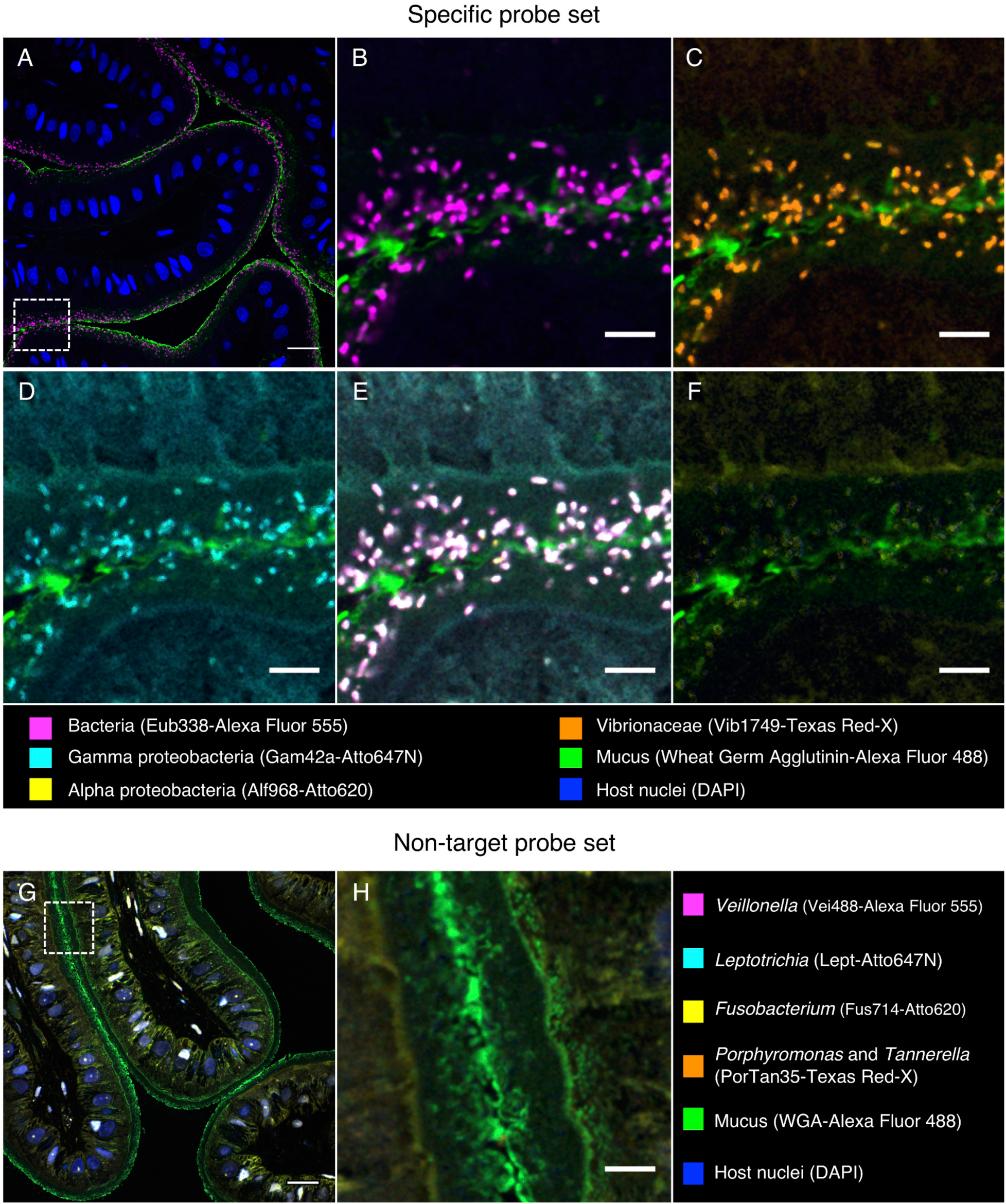
Fluorescence *in situ* hybridization identifies bacteria in the esophagus of *S. officinalis* as Vibrionaceae. A methacrylate-embedded section was hybridized with a nested probe set containing probes for most bacteria, Gammaproteobacteria, Alphaproteobacteria, and Vibrionaceae. (A) near-universal probe showing a similar bacterial distribution as in Figure 3. (B, C, D) Enlarged images of the region marked by the dashed square in (A) showing hybridization with near-universal, Vibrionaceae, and Gammaproteobacteria probes, respectively. (E) Merged image of B, C, and D showing an exact match of the signal from those three probes. (F) Alphaproteobacteria probe showing no hybridization. (G) An independent hybridization with the non-target probe set as a control; no signal is observed. (H) enlarged image of the dashed square in (G). Scale bar=20 µm (A, G); 5 µm (B-F, H).

**Table 2.**
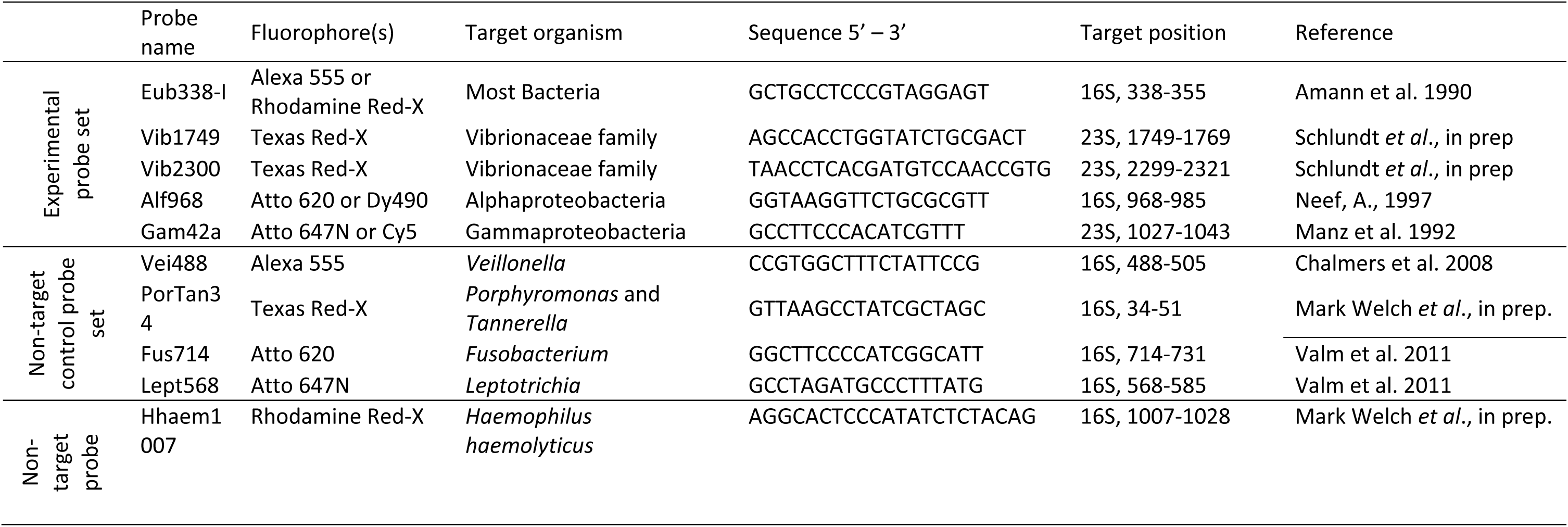
FISH probes used in this study

In other parts of the digestive tract we observed a sparser distribution of bacteria without obvious spatial organization. Bacteria in the intestine were present not in a layer but scattered throughout the lumen and mixed with the luminal contents (Fig. 5). Similarly, in the cecum, we observed bacteria in low abundance in the lumen (Fig. 6). FISH was also applied to the stomach, posterior salivary gland (poison gland) and duct of the salivary gland, but no bacteria were detected (not shown).

**Figure 5.**
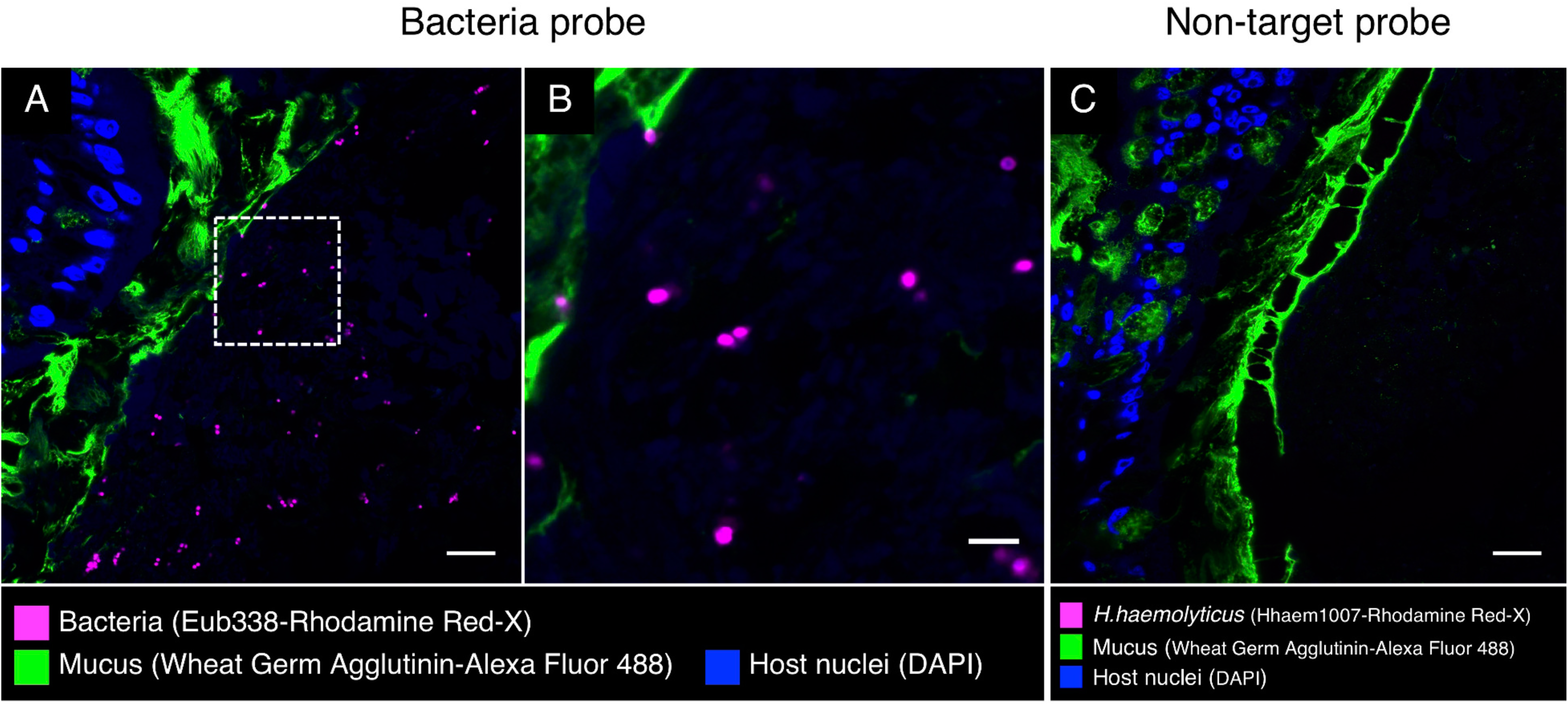
Fluorescence *in situ* hybridization in intestine of *S. officinalis*. A methacrylate-embedded section was hybridized with the near-universal probe and stained with fluorophore-labeled wheat germ agglutinin to visualize mucus. (A) Bacteria (magenta) are sparsely distributed through the lumen. (B) Enlarged image of the dashed square in (A). (C) An independent FISH control with a non-target probe (Hhaem1007); no signal was detected. Scale bar= 20 µm (A, C); 5 µm (B).

**Figure 6.**
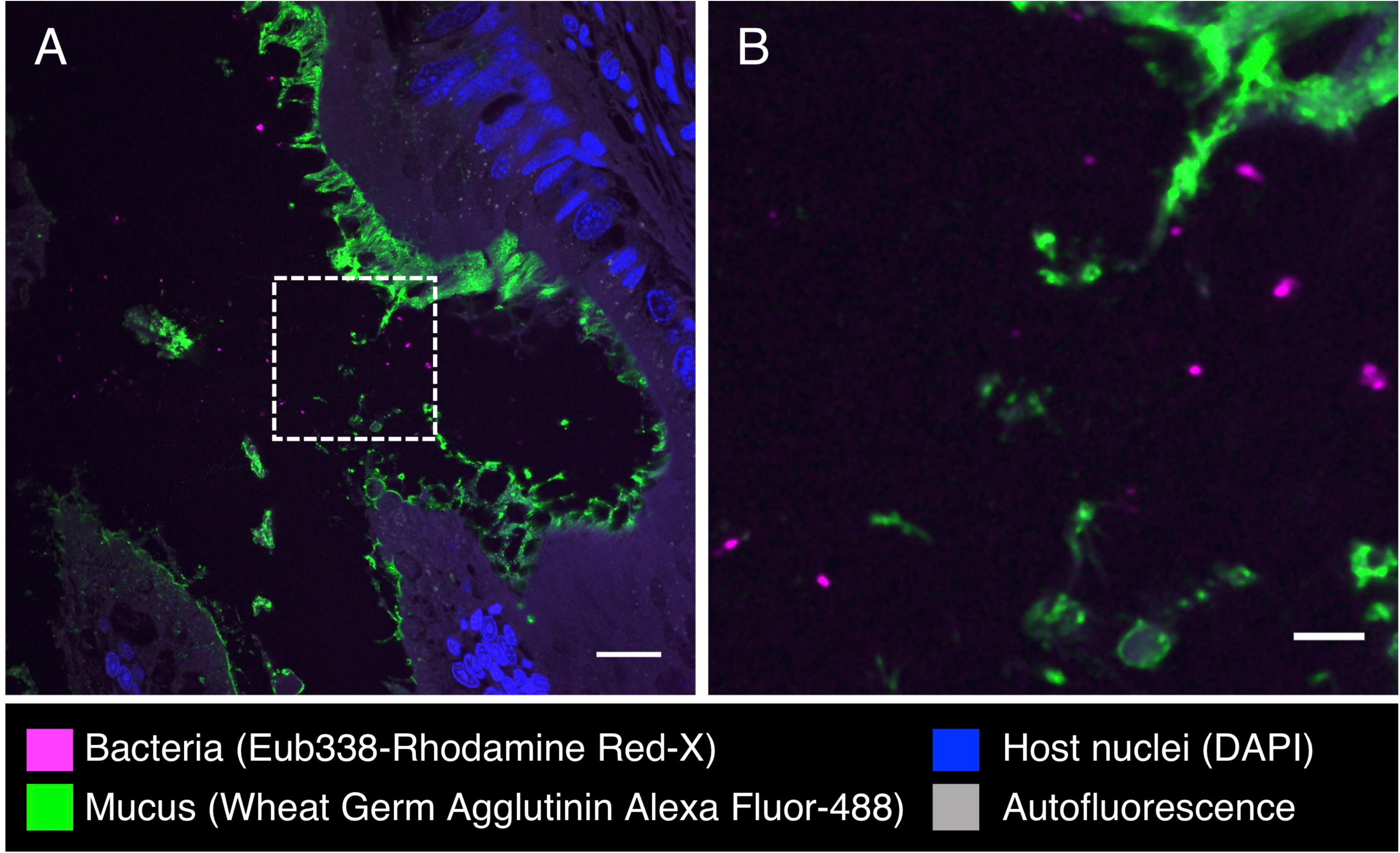
Fluorescence *in situ* hybridization in cecum of *S. officinalis*. (A) Bacteria are observed in low abundance in the lumen of cecum. (B) Enlarged image of dashed square in (A). Scale bar=20 µm (A); 5 µm (B).

Fluorescence *in situ* hybridization to cross-sections of the gills revealed clusters of bacteria at or near the surface of the tissue (Fig. 7). These bacteria hybridized with the Eub338 near-universal probe and a probe for Gammaproteobacteria (Fig. 7B, C, Table 2) but not Alphaproteobacteria (not shown), consistent with the identification of the clusters of gill bacteria as members of the gammaproteobacterial family Piscirickettsiaceae.

**Figure 7.**
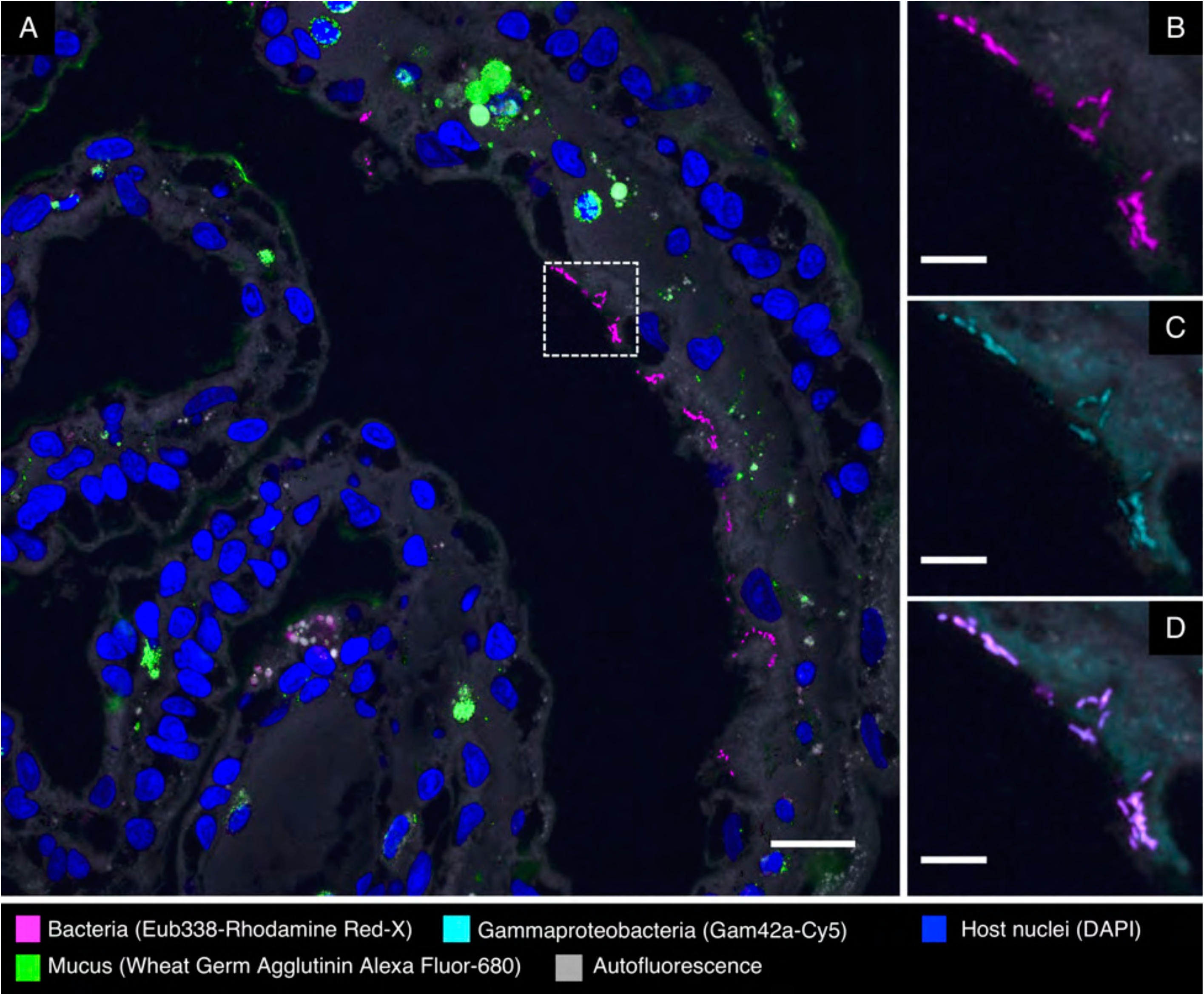
Fluorescence *in situ* hybridization in gills of *S. officinalis*. Bacteria are observed in small clusters at or near the surface of the gill. (A) Overview image. (B, C): enlarged images of the dashed square in (A) showing hybridization with near-universal and Gammaproteobacteria probes, respectively. (D) Merged image of (B) and (C) showing colocalization of the signal from those two probes. Scale bar=20 µm (A); 5 µm (B-D).

## 3. DISCUSSION

We sampled the cuttlefish microbiome of the digestive tract, gills, and skin and found dense colonization of an unexpected site, the esophagus, by a bacterium of the genus *Vibrio*. Both imaging and 16S rRNA gene sequencing showed a near-monoculture of *Vibrionaceae* in the esophagus, with imaging showing dense colonization of the interior lining of the esophagus with a single morphotype that hybridized to probes targeting *Vibrionaceae*. In the remainder of the GI tract, both imaging and 16S rRNA sequencing indicated a less consistent microbiota. Sequencing also showed lower relative abundance of the dominant *Vibrio* ASV, and qPCR confirmed a significantly lower total abundance of *Vibrio* cell copies in the distal GI tract. Imaging revealed sparse and sporadic colonization in the intestine and cecum, with scattered cells in the lumen and no clear colonization of the epithelium. In light of these results, we conclude that the GI tract of laboratory cultured *Sepia officinalis* has a highly simplified microbiome dominated by the genus *Vibrio*.

Diverse associations with *Vibrio* and the Vibrionaceae are known from cephalopods. Among the most extensively investigated is the mutualistic association of the bioluminescent *Vibrio (Aliivibrio) fischeri* with the light organ of the bobtail squid *Euprymna scolopes* (1, 13). Other well-known symbioses include the colonization of the cephalopod accessory nidamental gland with Alpha- and Gammaproteobacteria, which enables the host to secrete a layer of bacteria into the protective coating of the egg capsule (3, 14–18). Thus, colonization by Gammaproteobacteria and specifically by Vibrionaceae is common in cephalopods, yet colonization of the GI tract, and particularly the esophagus, was unexpected.

Bacteria from genus *Vibrio* and the related Vibrionaceae genus *Photobacterium* are frequent colonizers of the GI tracts of marine fishes (19, 20) and are prominent in the gastrointestinal microbiota of *Octopus vulgaris* paralarvae (21). Vibrionaceae have been reported to produce chitinases, proteases, amylase, and lipase (20), suggesting the possibility that colonization of the digestive tract by the Vibrionaceae serves to aid in host digestion (20). If the *Vibrio* and *Photobacterium* ASVs serve this function, their localization in high density in the esophagus, near the beginning of the digestive tract, may serve to seed the distal gut; colonization of the lining of the esophagus may provide a reservoir that permits the microbes to avoid washout from the gut by continually re-populating the lumen of downstream gut chambers.

An alternative explanation is that the colonization of the esophagus, and the rest of the gut, is pathogenic or opportunistic. Various *Vibrio* species are known pathogens of cephalopods, causing skin lesions and sometimes mortality in squids and octopuses (22–24). The genus *Vibrio* includes representative species that are pathogenic to corals (*V. coralliilyticus*), fish (*V. salmonicida*), diverse marine organisms (*V. harveyi*) and humans (*V. alginolyticus*, *V. cholerae*, *V. parahaemolyticus*, and *V. vulnificus)* (25, 26). Likewise, the genus *Photobacterium* contains pathogenic as well as commensal representatives (27). A previous study of the microbiota of *Octopus vulgaris* paralarvae found that recently hatched paralarvae had a high-diversity microbiome that changed, in captivity, to a lower-diversity microbiome with abundant Vibrionaceae (21). Whether the Vibrionaceae are an integral part of cuttlefish physiology in nature or whether they represent opportunistic colonists of these laboratory-reared organisms is a question for future research.

Our sequence data from gills were dominated by a single ASV classified as Piscirickettsiaceae that was in low abundance at other body sites. The Piscirickettsiaceae are a family within the Gammaproteobacteria (28) that includes the salmon pathogen *P. salmonis*. Rickettsia-like organisms have been described from the gills of clams and oysters (29, 30) as well as associated with the copepod *Calanus finmarchicus* (31). In recent years Piscirickettsiaceae have been identified in high-throughput sequencing datasets from seawater and sediment as a taxon that may be involved in biodegradation of oil and other compounds (32–38). Whether taxa in this family colonize the gills of cuttlefish and other organisms as symbionts or as opportunistic pathogens is a subject for future investigation.

Studies of wild *S. officinalis* microbiota will be informative for understanding natural host-symbiont associations under natural conditions, as compared to the mariculture-reared animals in the present study. *S. officinalis* in the eastern Atlantic and Mediterranean are known to prey on small mysids (crustaceans) in their first few weeks post-hatching; then as juveniles and adults they prey mainly on marine fishes and crabs. The sparseness and simplicity of the gut microbiota observed in our study may have been in part a result of the monodiet of grass shrimp (*Palaemonetes*) we employed. It remains to be seen whether differences in diet and natural variation in environmental conditions influence the association of microbial symbionts with *S. officinalis* in the wild.

Because cuttlefish behavior is well-studied and there exist standardized methods for documenting multiple behaviors (8), we hypothesized that these animals may provide a unique opportunity to study microbes and the gut-brain axis – the effect of gut microbiota on behaviour (39) – in an invertebrate system. Therefore, in parallel with our study of the microbiome of various organ systems, we conducted extensive preliminary experiments to study the effect of antibiotic treatment on the behavior of *S. officinalis*. These results were largely negative, perhaps due to the highly simplified microbiota we observed and its resilience to the antibiotic employed. These results may prove helpful to the cephalopod husbandry community, as they suggest that application of the commonly used antibiotic enrofloxacin is compatible with maintenance of normal behavioral and microbiome characteristics of this species

## 4. MATERIALS AND METHODS

### 4.1 Sampling and antibiotic treatment

Our study included 27 cuttlefish that were bred in the wild and moved to captivity prior to hatching. Animals were held in water tables connected to a single open-filtration system fed by filtered seawater. Animals were euthanized via immersion into a 10% dilution of ethanol in seawater and were then dissected under sterile conditions within a biosafety cabinet using autoclaved tools to obtain samples for microbial analyses. The gastrointestinal tract was dissected into four components: esophagus, stomach, cecum, and intestine. Gill tissue and skin from the mantle was sampled as well (∼ 0.5g per sample). All tissues were stored in separate sterile cryogenic tubes and flash-frozen in liquid nitrogen.

Following a pilot study of three individual cuttlefish, we included 24 cuttlefish (16 test, 8 control) in an experiment designed to test the effect of the antibiotic treatment on the composition of the cuttlefish microbiome. The experimental design consisted of administering antibiotic to animals in the treatment group (n = 16) via injection into the food source (grass shrimp, *Palaemonetes* sp.), which was then fed to the animals. Prior to feeding, shrimp were injected with enrofloxacin (Baytril^®^; 22.7 mg/mL, Bayer HealthCare LLC, Shawnee Mission, KS, USA) using a 0.5 cc, U-100 insulin syringe with an attached 28 g x 1/2” needle (Covidien LLC, Mansfield, MA, USA). The antibiotic dosage was 10 mg/kg rounded up to the nearest hundredth mL. The antibiotic was injected into the coelomic cavity of the shrimp which were then immediately fed to the cuttlefish once daily for 14 days. We maintained 8 animals as controls, none of which received antibiotic treatment. Experimental animals were held in two separate water tables and control animals a third, all of which were connected to the same open-filtration system fed by filtered seawater. Within each water table, animals were isolated into individual holding pens via plastic panels. The experimental period lasted for 14 days (1 – 15 Oct 2017), during which fecal samples were collected from each individual daily; fecal samples were collected via pipetting free-floating material from the tank of each individual animal into a sterile pipette.

To assess the extent to which cuttlefish microbial symbionts were shared with their environment and food sources, 1L water samples were taken from each of three water tables in which animals were held on the day of euthanasia and filtered using a 0.22 micron Sterivex filter for DNA extraction; grass shrimp used as the food source throughout the duration of the experiment were collected in 1.8ml sterile cryotubes on the same day that water was sampled, and were frozen at −20°C until prepared for DNA extraction.

### 4.2 DNA extraction, sequencing, qPCR, and16S rRNA gene statistical analyses

DNA extractions were performed on cuttlefish tissue biopsies, water, and whole shrimp using the MoBio PowerSoil 96 Well Soil DNA Isolation Kit (Catalog No. 12955-4, MoBio, Carlsbad, CA, USA). We used the standard 515f and 806r primers (49–51) to amplify the V4 region of the 16S rRNA gene, using mitochondrial blockers to reduce amplification of host mitochondrial DNA. Sequencing was performed using paired-end 150 base reads on an Illumina HiSeq sequencing platform. Following standard demultiplexing and quality filtering using the Quantitative Insights Into Microbial Ecology pipeline (QIIME2) (52) and vsearch8.1 (53), ASVs were identified using the Deblur method (19) and taxonomy was assigned using the Greengenes Database (May 2013 release; http://greengenes.lbl.gov). Libraries containing fewer than 1000 reads were removed from further analyses. All 16S rRNA sequence data and metadata will be made available via the QIITA platform prior to publication. Statistical analyses and figure production were produced with the aid of several R packages including vegan (40), dplyr (41), ggplot2 (42) and Adobe Illustrator® CC 2019. Code for analyses and figures is available at www.github.com/hollylutz/CuttlefishMP. We performed qPCR analyses targeting the genus *Vibrio* to quantify the abundance of bacterial cells throughout the GI tract. Conditions for qPCR were borrowed from Thompson et al. (43), using the primers 567F and 680R and performing two replicates of qPCR analysis that were combined to produce the final results reported in this study.

### 4.3 Sample collection, fixation and sectioning for imaging

Samples from esophagus, stomach, intestine and cecum of 9 cuttlefish (1 from the pilot and 8 from the second period experiment) were dissected and divided in order to include the same individuals in both microscopy and sequencing analyses. Immediately after dividing, samples for imaging were fixed with 2% paraformaldehyde in 10 mM Tris pH 7.5 for 12 h at 4 °C, washed in PBS, dehydrated through an ethanol series from 30, 50, 70, 80 and 100%, and stored at −20 °C. Samples were dehydrated with acetone for 1 h, infiltrated with Technovit 8100 glycol methacrylate (EMSdiasum.com) infiltration solution 3 times 1 hour each followed by a final infiltration overnight under vacuum, transferred to Technovit 8100 embedding solution and solidified for 12 h at 4 °C. Blocks were sectioned to 5 um thickness and applied to Ultrastick slides (Thermo Scientific). Sections were stored at room temperature until FISH was performed.

### 4.4 Fluorescence *in situ* hybridization (FISH)

Hybridization solution [900 mM NaCl, 20 mM Tris, pH 7.5, 0.01% SDS, 20% (vol/vol) formamide, each probe at a final concentration of 2 μM] was applied to sections and incubated at 46 °C for 2 h in a chamber humidified with 20% (vol/vol) formamide. Slides were washed in wash buffer (215 mM NaCl, 20 mM Tris, pH 7.5, 5mM EDTA) at 48 °C for 15 min. Samples were incubated with wheat germ agglutinin (20 ug ml^−1^) conjugated with Alexa Fluor 488 and DAPI (1 ug ml^−1^) at room temperature for 30 minutes after FISH hybridization to label mucus and host nuclei, respectively. Slides were dipped in ice-cold deionized water, air dried, mounted in ProLong Gold antifade reagent (Invitrogen) with a #1.5 coverslip, and cured overnight in the dark at room temperature before imaging. Probes used in this study are listed in Table 2.

### 4.5 Image acquisition and linear unmixing

Spectral images were acquired using a Carl Zeiss LSM 780 confocal microscope with a Plan-Apochromat 40X, 1.4 N.A. objective. Images were captured using simultaneous excitation with 405, 488, 561, and 633 nm laser lines. Linear unmixing was performed using the Zeiss ZEN Black software (Carl Zeiss) using reference spectra acquired from cultured cells hybridized with Eub338 probe labeled with the appropriate fluorophore and imaged as above. Unmixed images were assembled and false-colored using FIJI software (Schindelin *et al.*, 2012).

### 4.6 Data availability

All 16S rRNA sequences and sample metadata will be made available via the QIITA platform prior to publication.

## ACKNOWLEDGEMENTS

We thank Alan Kuzirian, Louie Kerr, Wyatt Arnold, Madeline Kim, Elle Hill, Tyler He, Tifani Anton, and Eric Edsinger. HLL was supported by NSF 1611948. TR and JMW were supported by NSF 1650141. RTH thanks the Sholley Foundation for partial support. Amber Durand was supported by the Woods Hole Partnership Education Program. The funders had no role in study design, data collection and interpretation, or the decision to submit the work for publication.

